# Stochastic model of Alzheimer’s Disease progression using two-state Markov chains

**DOI:** 10.1101/2023.06.29.547071

**Authors:** Meaghan Parks

## Abstract

In 2016, Hao and Friedman developed a deterministic model of Alzheimer’s disease progression using a system of partial differential equations. This model describes the general behavior of the disease, however, it does not incorporate the molecular and cellular stochasticity intrinsic to the underlying disease processes. Here we extend the Hao and Friedman model by modeling each event in disease progression as a stochastic Markov process. This model identifies stochasticity in disease progression, as well as changes to the mean dynamics of key agents. We find that the pace of neuron death increases whereas the production of the two key measures of progression, Tau and Amyloid beta proteins, decelerates when stochasticity is incorporated into the model. These results suggest that the non-constant reactions and time-steps have a significant effect on the overall progression of the disease.

## Introduction

Alzheimer’s disease (AD) affects approximately 5.8 million people in the United States and is the most common cause of dementia (CDC 2020). The CDC further predicts 14 million instances of the disease in the US by 2060 (2020). Despite years of research into the disease, its progression is poorly understood and ineffectively treated.

Nevertheless, we know that the malfunction of certain proteins results in inflammation and neuron death. Two proteins in particular are suspected to drive disease progression: Amyloid Beta and Tau (Hao & Friedman 2016). Build up of Amyloid Beta forms plaques in the brain, which interfere with communication between cells and are toxic to neurons (Hu et al. 2020). Healthy tau proteins aid in neuronal transport and support neuron structure; however, in Alzheimer’s disease, collections of misfolded Tau proteins create toxic Neurofibrillary tangles (Hao & Friedman 2016).

Existing mathematical models of Alzheimer’s disease are either entirely deterministic, such as Hao & Friedman’s model (which uses Ordinary Differential Equations, ODEs (2016)], or incorporate stochastic noise in an *ad hoc* manner, e.g. as a separate variable as proposed by Zhang & Wang (2020). Here, we propose a stochastic model that represents every state variable in disease progress probabilistically, as a random variate in a Markov process. We leverage the same mechanistic model as proposed by Hao & Friedman (after consolidating a few parameters). Because there are many interacting molecules in the model, we hypothesize that variability will compound and alter mean behavior – as typically seen in complex systems with non-linear interactions between agents, e.g. cell proliferation or gene transcription (Potvin-Trottier et al. 2016; Yates et al. 2017). Furthermore, we predict that compounds that interact with more other molecules will have the highest variability, namely Amyloid Beta outside the neurons.

## Methods

### Overview

The model detailed in this paper is based upon the deterministic model proposed by Hao and Freidman (2016), with a few simplifying changes that do not appreciably alter dynamics in the deterministic case. To increase the interpretability of the stochastic model, the number of parameters was reduced from 77 to 29, and the number of state variables was reduced from 11 to 9. These changes leave only parameters/variables thought to be most important to the disease’s progression (discussed below). For direct comparison between the deterministic and stochastic models, we implemented a deterministic version of the model with the same reduction in parameters and state variables.

### Variables & Parameters

The nine variables on which the model relies are shown in Table 1, the table also displays the initial values used for both models. The variables used in the model were chosen based on the prevailing theory that the driving force behind Alzheimer’s Disease is the formation of Amyloid plaques and Neurofibrillary tangles.

**Table 1.** Initial Values

| Variable | Description | Value |
| --- | --- | --- |
| $A_{\beta}^i$ | Amyloid Beta made inside neurons | $10^{-6}g/ml$ |
| $A_{\beta}^o$ | Amyloid Beta outside neurons | $10^{-8}g/ml$ |
| $\tau$ | Tau Protein | $1.37 \times 10^{-10}g/ml$ |
| $F_i$ | NFTs inside neurons | $3.36 \times 10^{-10}g/ml$ |
| $F_o$ | NFTs outside neurons | $3.36 \times 10^{-11}g/ml$ |
| A | Astroglia | $.14g/ml$ |
| M1 | Proinflammatory Microglia | $.02g/ml$ |
| M2 | Anti-inflammatory Microglia | $.02g/ml$ |
| N | Neurons | $.14g/ml$ |

The models use twenty-nine parameters displayed in Table 2; asterisks identify parameters from Hao & Friedman’s model (2016). Only four parameters (no asterisks, Table 2) were modified in the newly proposed model. The parameter □_Nd_ represents the rate of normal neuronal loss with age; after the age of forty, neurons begin to die at ∼5% each year. The parameter □MA is a combination of two separate parameters used by Hao & Friedman: the production of TNF-alpha inflammatory microglia and the production of astrocytes by TNF-alpha. Because we omitted TNF-alpha in the model, we multiplied the two values to bridge the gap between the astrocytes and inflammatory microglia, both of which are elements of our model. □_M1_ and □_M2_ represent the effect of inflammation on inflammatory microglia □_M1_ and its effect on anti-inflammatory microglia □_M2_.

**Table 2.** Parameters

| Parameter | Description | Value |
| --- | --- | --- |
| $\lambda_{Nd}$ | Neuron death rate | $\frac{.05}{365}/day$ |
| $\lambda_{\beta}^i$ | Production rate of $A_{\beta}^i$ | $9.51 \times 10^{-6} g/ml/day$ * |
| $\lambda_N$ | Production rate of $A_{\beta}^o$ by neuron | $8 \times 10^{-9} g/ml/day$ * |
| $\lambda_{MA}$ | Production rate of astrocytes by M1 | $.045 g/ml/day$ |
| $\lambda_A$ | Production rate of $A_{\beta}^o$ by astrocytes | $8 \times 10^{-10} g/ml/day$ * |
| $\lambda_{\tau}$ | Production rate of Tau proteins by ROS | $1.35 \times 10^{-11} g/ml$ * |
| $\lambda_F$ | Production rate of NFT by Tau | $1.662 \times 10^{-3}/day$ * |
| $\lambda_{AA_{\beta}^o}$ | Production/Activation rate of astrocytes by $A_{\beta}^o$ | $1.793/day$ * |
| $\lambda_{\tau_0}$ | Production of Tau proteins (healthy) | $8.1 \times 10^{-11}/day$ * |
| $\lambda_{MF}$ | Production/Activation rate of Microglia by NFT | $2 \times 10^{-2}/day$ * |
| $d_{Fi}$ | Degradation rate for $F_i$ | $2.77 \times 10^{-3}/day$ * |
| $d_{F_0}$ | Degradation rate for $F_i$ | $2.77 \times 10^{-4}/day$ * |
| $d_{M1}$ | Degradation rate for M1 | $.015/day$ * |
| $d_{M2}$ | Degradation rate for M2 | $.015/day$ * |
| $d_{A_{\beta}^o_M}$ | Clearance rate of $A_{\beta}^o$ by microglia | $2 \times 10^{-3}/day$ * |
| $d_{A_{\beta}^i}$ | Clearance rate of $A_{\beta}^i$ | $9.51/day$ * |
| $d_A$ | Death rate of astrocytes | $1.2 \times 10^{-3}/day$ * |
| $d_{NF}$ | Death rate of neurons by NFT | $3.4 \times 10^{-4}/day$ * |
| $d_{\tau}$ | Degradation of $\tau$ proteins | $0.277/day$ * |
| $K_{Fi}$ | Half-saturation of intracellular NFTs | $3.36 \times 10^{-10} g/ml$ * |
| $K_{Fo}$ | Average of extracellular NFTs | $2.58 \times 10^{-11} g/ml$ * |
| $K_{A_{\beta}^o}$ | Michaelis-Menton coefficient for $A_{\beta}^o$ | $7 \times 10^{-3} g/cm^3$ * |
| $M_G^0$ | Source of Microglia | $.047 g/cm^3$ * |
| $\beta_{M1}$ | Measure of inflammation | .9 |
| $\beta_{M2}$ | Measure of inflammation | .09 |
| $\theta$ | M1/M2 effectivity in clearance of $A_{\beta}^o$ | .9 * |
| $R_0$ | Initial inflammation by ROS | 6 * |
| $N_0$ | Reference density of neurons | $.14 g/cm^3$ * |
| $A_0$ | Reference density of astrocytes | $.14 g/cm^3$ * |

### Deterministic Model

The deterministic model we created, to which we will compare the stochastic, is represented by the compartment model in Figure 1 and modeled as a system of 10 ODEs (Table 3).

**Table 3:**
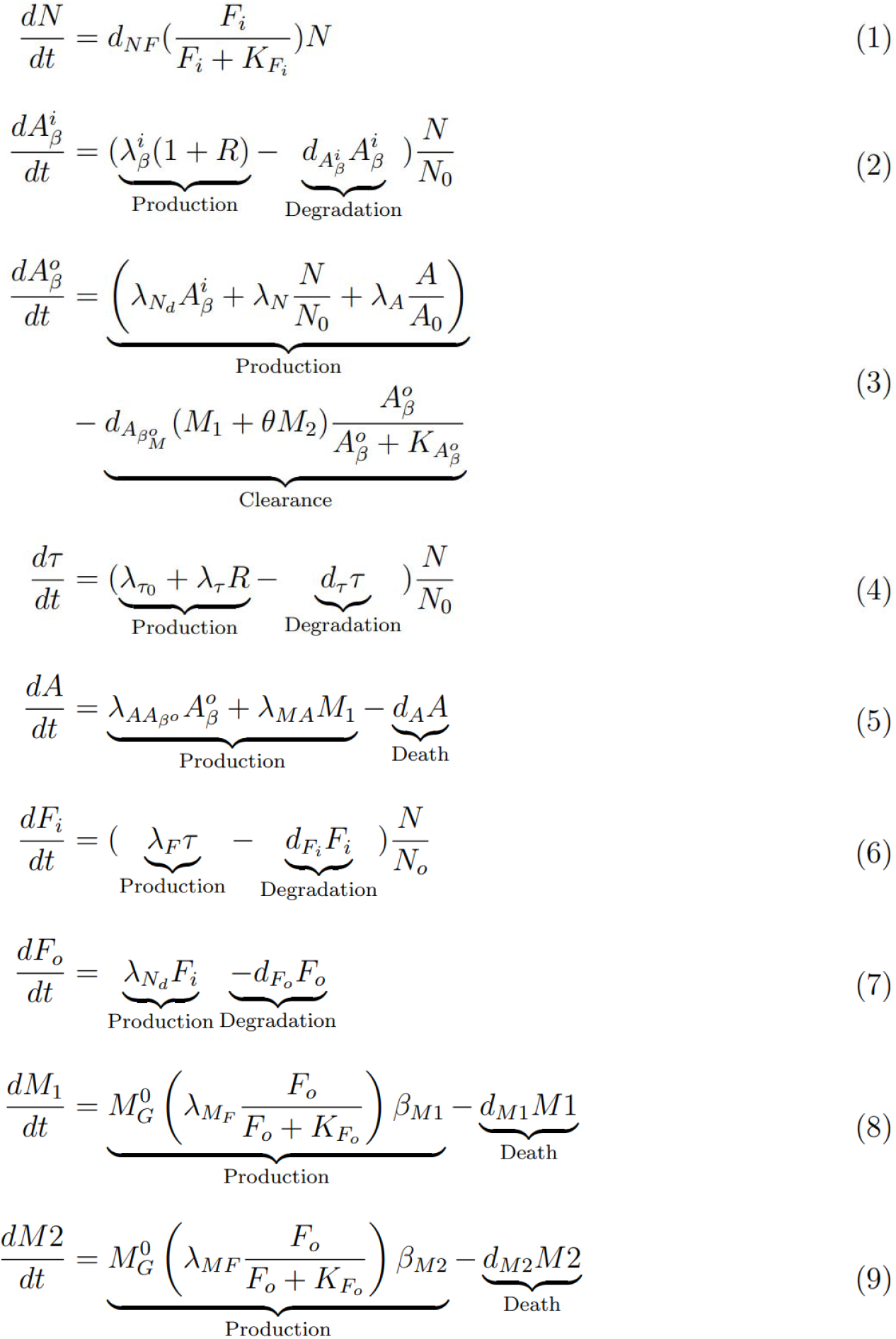
Deterministic Model Equations

**Figure 1.**
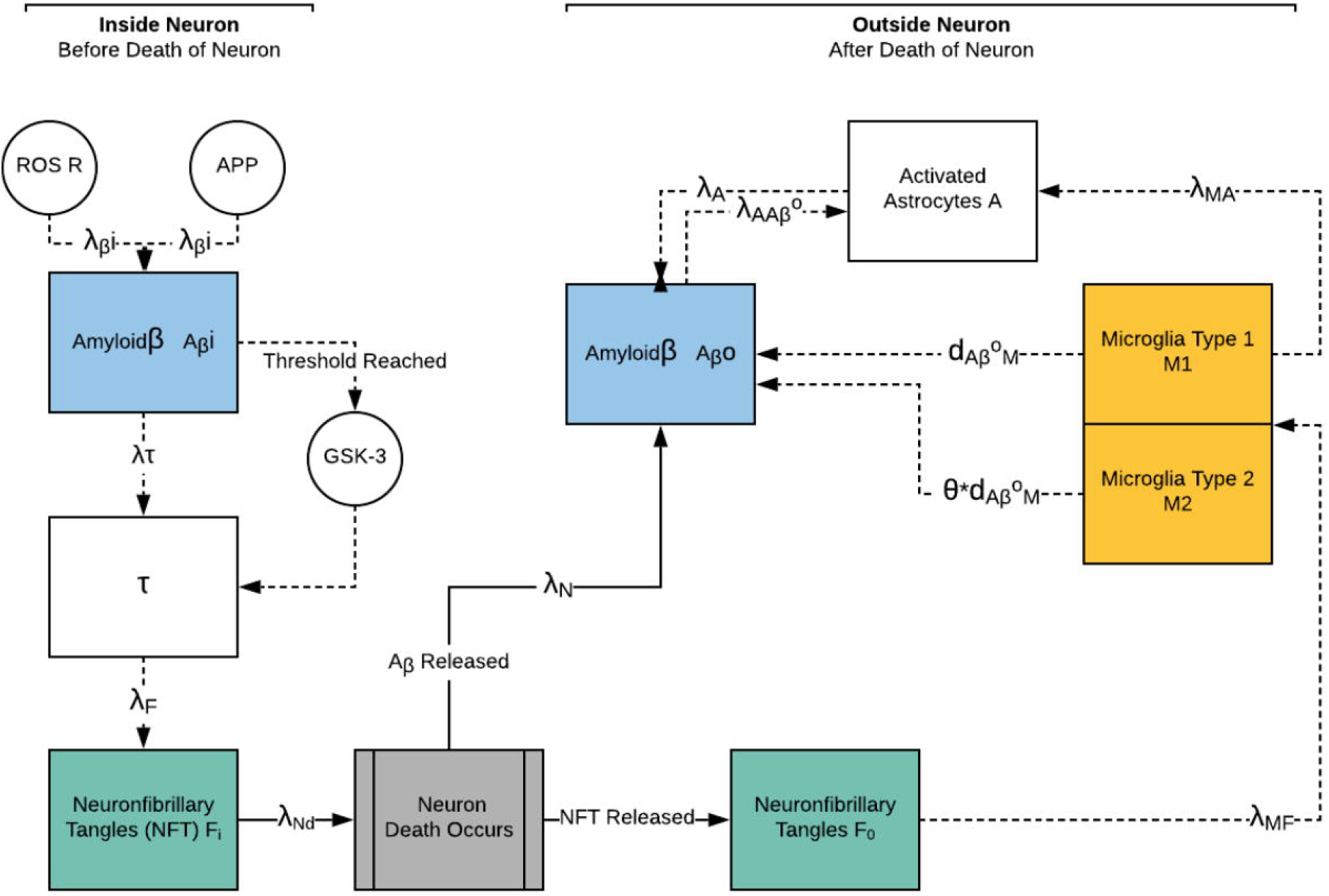
Compartment Model

Thematically, in short:

- Equation 1 is the change in neuron concentration; only living neurons divide.
- Equation 2: change in the concentration of Amyloid Beta inside neurons, whereas,
- Equation 3 is the concentration of Amyloid Beta that the neuron releases when it dies.
- Equation 4: change in the concentration of Tau proteins;
- Equation 5 is the change in concentration of Astrocytes.
- Equations 6 and 7 are analogous to equations 2 and 3 but for Neurofibrillary tangles rather than Amyloid Beta.
- Equations 8 and 9 represent the change in inflammatory and anti-inflammatory microglia concentrations, respectively.

The model was run for 10 years, corresponding to 3650 time-steps (365 days x 10 years), which is the average life expectancy for a person with Alzheimer’s disease, with the initial values shown in Table 1 and the parameters in Table 2. Because of the deterministic nature of the model, the model yields identical results every iteration.

### Stochastic Model

We then converted the deterministic model to a stochastic model by conceptualizing each variable as a two-state Markov process. Table 4 describes each variable, the two associated states, and the processes by which quanta enter and leave the states. The stoichiometry matrix, which contains the changes in the molecules concentration for each reaction, contains the equations from the deterministic model. Therefore, the stoichiometry matrix for the stochastic model is not constant and the amount by which the concentration of a molecule changes depends on the concentration of the other relevant molecules.

**Table 4.** Markov States and Transitions

| Variable | State 1 | State 2 | Process(es) |
| --- | --- | --- | --- |
| $A^i_\beta$ | Inside the neuron | Outside the neuron | Birth, Emmigration |
| $A^o_\beta$ | Outside the neuron | Degraded | Birth, Death |
| □ | Normal | Tangle | Birth, Death |
| $F_i$ | Inside neuron | Degraded/Released | Birth, Death, Emmigration |
| $F_o$ | Outside neuron | Degraded | Birth, Death |
| A | Normal | Degraded | Birth, Death |
| M1 | Normal | Degraded | Birth, Death |
| M2 | Normal | Degraded | Birth, Death |
| N | Normal | Degraded | Pure Death |

We then simulated activity using a second-order Gillespie algorithm. Functions were then created for each birth/death process. The functions determine whether or not the reactions will occur based upon the reaction rate relative to all the other reactions. For example, the function for neuron death draws a normally-distributed number *n*, with a mean equal to dNF over KFi divided by the sum of all the reaction rates. The normal distribution was used because the rates are averages. This number is then compared to a uniformly-distributed random variable *U ∼* [0, 1), determined using the runif() function, and, if *U* < *n*, the reaction will occur.

In terms of variables that have both production and degradation, either both reactions will occur, or no reaction will take place. For instance, amyloid beta inside the neuron has both production and degradation reactions. Again, a uniformly distributed random variable *U*, is compared with a normally distributed variable *n*, with a mean equal to the sum of the rate of production and rate of degradation divided by the sum of all reaction rates. In this case, if *U < n*, then both degradation and production reactions occur.

The time steps of the model varied in size and were determined in the Gillespie algorithm. The largest time was used to increment the count, however the time till the next reaction for each molecule was determined and used to determine the order in which the functions should be checked. In other words if the time until the next reaction for neurons was shorter than that of Amyloid Beta inside, the algorithm would first update the neuron value if necessary then Amyloid Beta.

The model is run for 10 years, the same as the deterministic model. However, because of the varying time step size in the stochastic model, different reactions may occur more or less than the deterministic model.

## Results

We tracked each of the nine variables over 25 iterations, and compared the stochastic and deterministic results for each below. Finally, the variance for each variable in the stochastic model was compared to test the above hypothesis that Tau and Amyloid Beta will have the highest variability. Summarily, all variables in the stochastic model progress in the same direction as predicted from the deterministic model (e.g. if the variable monotonically increases in the deterministic model, it does so in the stochastic model as well, Supplementary Figures 2-18). However, the rate of change of many variables changed dramatically in the stochastic model, and several instances of non-monotonic behavior disappeared.

**Figure 2.**
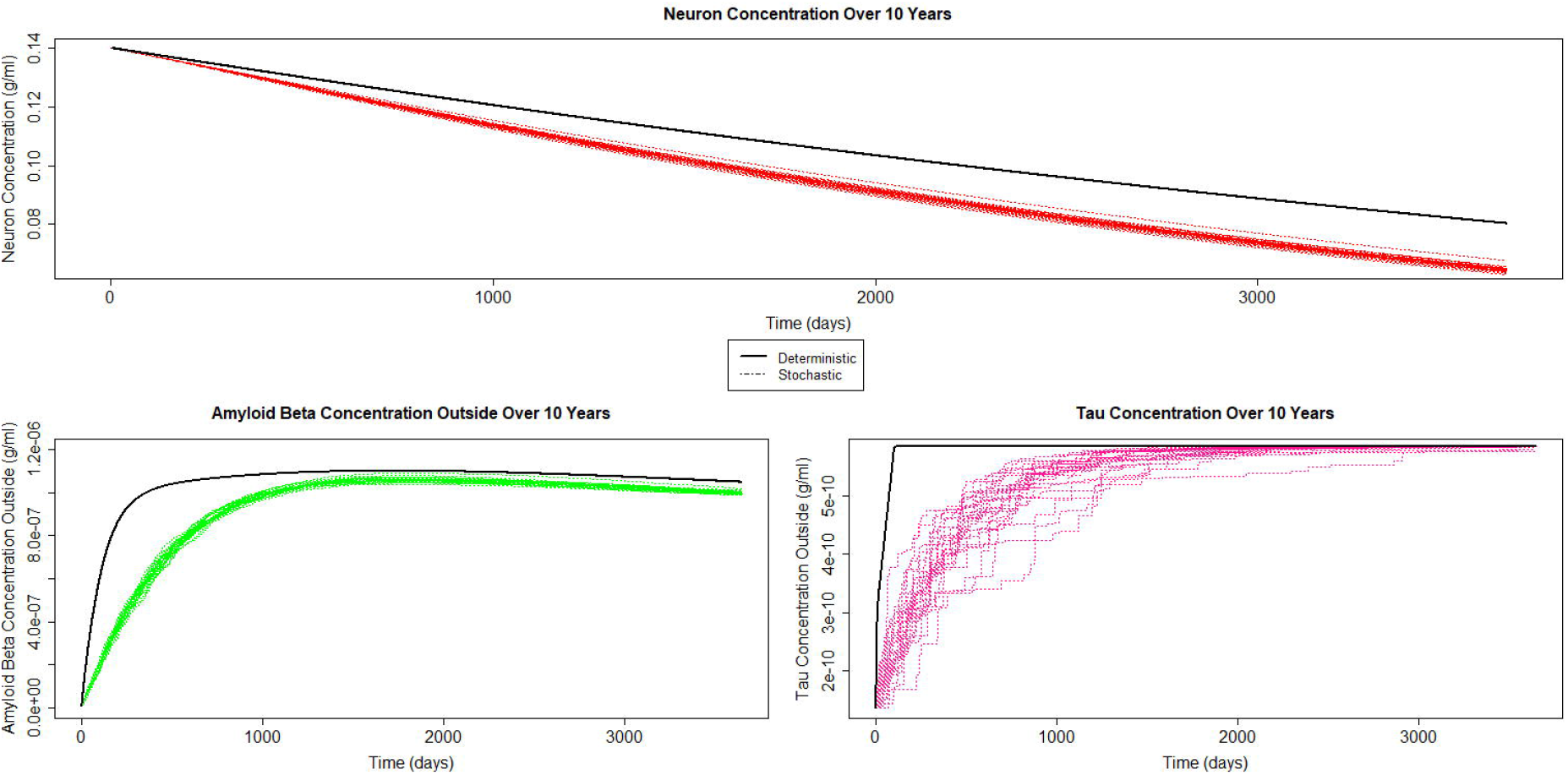
A) Neuron concentration over ten years, deterministic and stochastic. B) Amyloid Beta outside concentration over ten years, deterministic and stochastic. C) Tau concentration over ten years, deterministic and stochastic.

In the main text, we discuss dynamics of three key state variables: neuron concentration (Figure 2A), Amyloid beta outside the neuron (Figure 2B), and Tau (Figure 2C).

### Neurons

Figure 2A shows the results from the stochastic model for neurons from the stochastic model and from the deterministic model, with the dashed black line representing the latter. Comparing the output from both models shows that neurons die slower in the deterministic model.

### Amyloid Beta Outside

Amyloid Beta outside the neuron is the first reaction we will look at which has both production and degradation. The concentration of amyloid beta outside the neuron rises quickly before slowly decreasing. Figure 2B shows the output from stochastic and deterministic models for amyloid beta outside the neuron. The models have a similar shape, however the deterministic model, shown by the solid black line, increases faster than the stochastic model. Both models seem to begin transitioning from increasing to slowly decreasing at approximately the same time.

### Tau

Tau concentration in the stochastic model increases at a rate similar to that of amyloid beta outside the neuron. A crucial difference between the two is that tau reaches an equilibrium and stabilizes, whereas amyloid beta outside the neuron does not. The behavior of tau in the stochastic model can be seen in Figure 2C, which shows 25 different runs of the model.

Figure 2C also demonstrates the different behavior of tau in the stochastic and deterministic models. Tau concentration in the deterministic model increases appreciably faster than in the stochastic model, however despite this differing behavior both models do reach the same equilibrium.

## Other Variables

### Amyloid Beta Inside the Neuron

Amyloid Beta inside the neurons increases quickly before rapidly reaching equilibrium (see supplementary Figure 1). The comparison between the deterministic model and the stochastic model results for amyloid beta inside the neuron show that both models reach the same equilibrium at approximately the same time despite the fact that the reactions could be occurring more frequently in the stochastic model (see supplementary Figure 1).

### NFTs Inside the Neuron

The concentration of NFTs inside the neuron increases at a fast rate for approximately the first 1000 days, after which it begins to decrease (see supplementary Figure 2).

The difference between the deterministic and stochastic models are most stark when comparing NFT inside the neuron; this can be observed in Supplementary Figure 2 which shows 25 runs of the stochastic model compared with the deterministic model. In the deterministic model NFTs inside the neuron decrease sharply before increasing and leveling off, this is a significant departure from the behavior seen in the stochastic model.

### NFTs Outside the Neuron

The concentration of NFTs outside the neuron increases over the 10 years, as shown in Supplementary Figure 3, this figure also shows that the concentration of NFTs outside the neuron increases faster in the deterministic model than in the stochastic model, despite NFTs inside the neuron increasing faster in the stochastic model.

**Figure 3.**
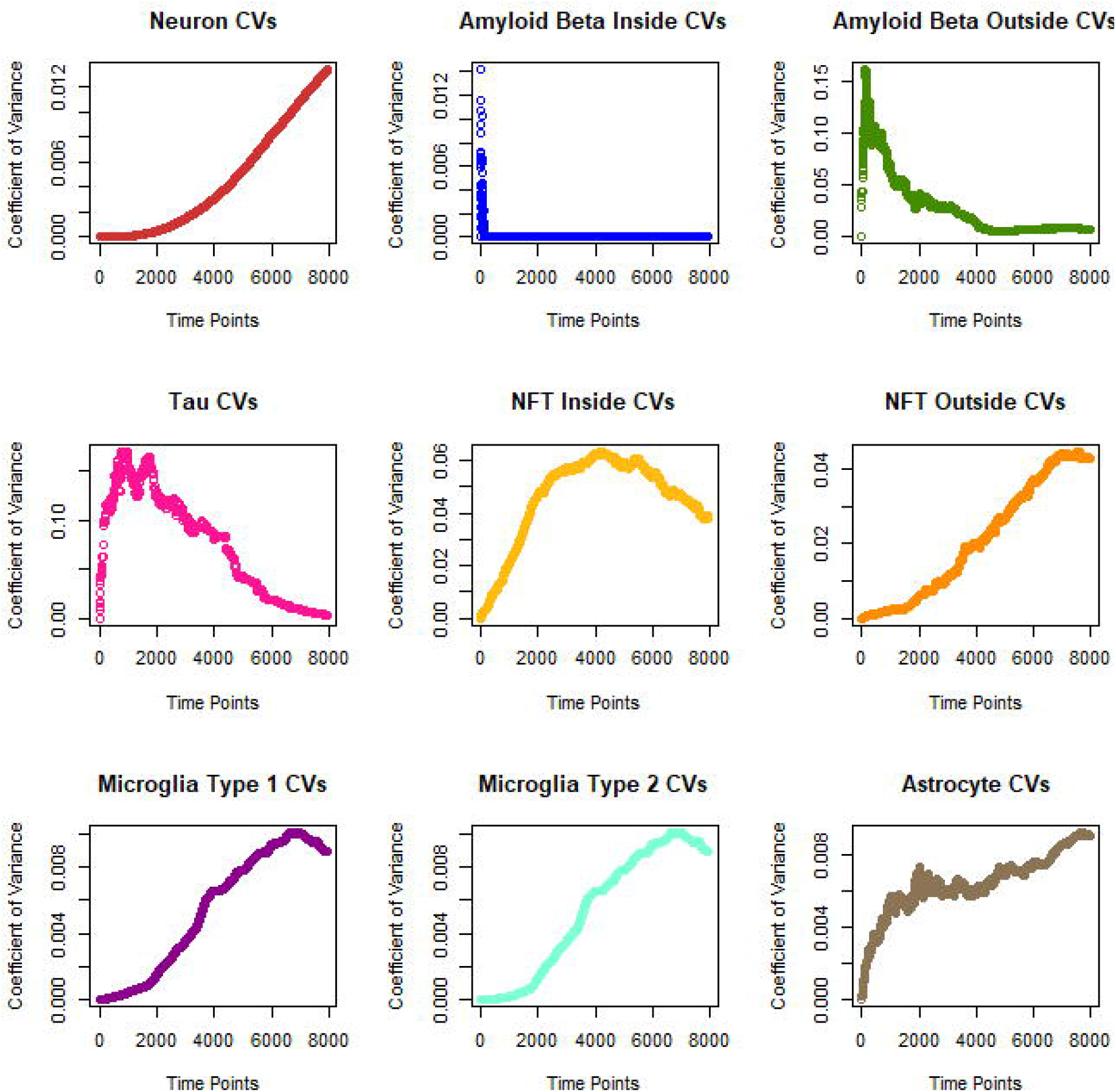
Change in CV overtime

### Immune Cells: Astrocytes & Microglia

Astrocyte concentration in the stochastic model increases over time with the rate of increases slowing the longer the model runs, Supplementary Figure 4A shows this behavior in 25 runs of the stochastic model. Astrocytes concentration in the deterministic model behaves in a similar manner however the concentration increases slightly faster than in the stochastic model, also shown in supplementary Figure 4A. The faster increase in the deterministic model is logical as astrocyte production is triggered by type 1 microglia production of which is triggered by NFTs outside the neuron, which as shown above also increases more rapidly in the deterministic model than the stochastic model.

Type 1 microglia initially increases in concentration rapidly before beginning to slow at approximately 200 days. Type 2 microglia behaves quite differently and decreases rapidly before leveling off. Panels B and C of supplementary figure 4 respectively show type 1 and type 2 microglia concentration from 25 runs of the model. The behavior of type 1 and type 2 microglia in the deterministic model is very similar to in the stochastic model, with the only difference being that type 1 microglia increases faster in the deterministic model.

### Variance

The hypothesis tested by this work was that the compound within the model that would be the most variable would be Amyloid Beta outside the neurons. The model was run 25 times with matching time jumps, forcing the times allowed for variance between runs at each time point to be found; from this, the average variance was found for each compound, and the results were used to assess the validity of the hypothesis.

Table 5 shows that amyloid beta is not the most variable between runs of the model. Instead Tau is the most variable, despite being one of the state variables that interacts with the fewest other variables, 1 compared with amyloid beta outside’s 5 variables, the number of interactions comes from the equations in Table 3. This strongly disagrees with the proposed hypothesis that state variables which interact with more other state variables will be more variable. Figure 20 shows the coefficient of variation (CV), which is the ratio of the standard deviation to the mean, at each time point, demonstrating which molecules reach equilibrium.

**Table 5.**
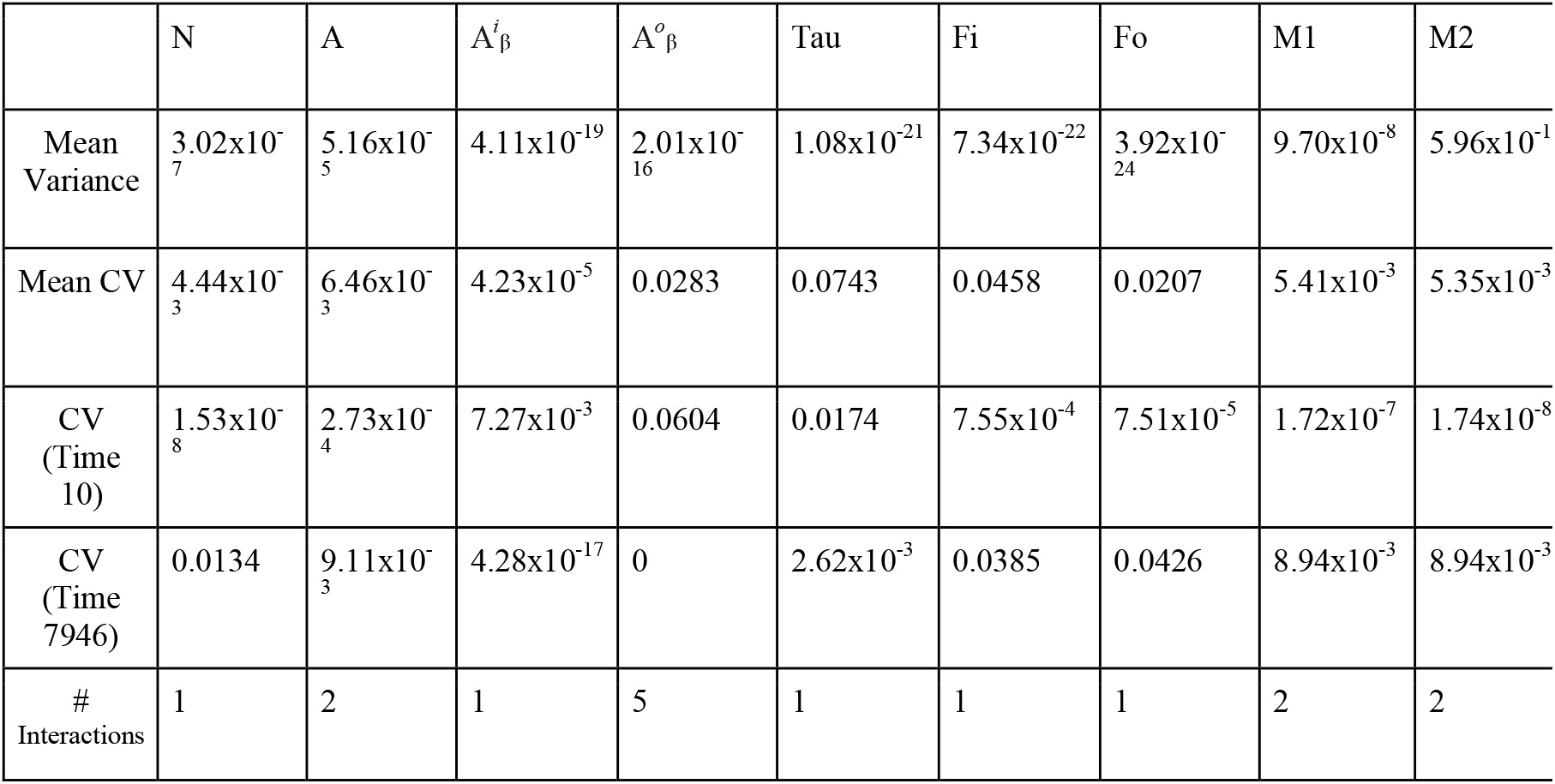
Variability of state variables

| | N | A | $A_{\beta}^i$ | $A_{\beta}^o$ | Tau | Fi | Fo | M1 | M2 |
| --- | --- | --- | --- | --- | --- | --- | --- | --- | --- |
| Mean<br>Variance | $3.02 \times 10^{-7}$ | $5.16 \times 10^{-5}$ | $4.11 \times 10^{-19}$ | $2.01 \times 10^{-16}$ | $1.08 \times 10^{-21}$ | $7.34 \times 10^{-22}$ | $3.92 \times 10^{-24}$ | $9.70 \times 10^{-8}$ | $5.96 \times 10^{-11}$ |
| Mean CV | $4.44 \times 10^{-3}$ | $6.46 \times 10^{-3}$ | $4.23 \times 10^{-5}$ | 0.0283 | 0.0743 | 0.0458 | 0.0207 | $5.41 \times 10^{-3}$ | $5.35 \times 10^{-3}$ |
| CV<br>(Time<br>10) | $1.53 \times 10^{-8}$ | $2.73 \times 10^{-4}$ | $7.27 \times 10^{-3}$ | 0.0604 | 0.0174 | $7.55 \times 10^{-4}$ | $7.51 \times 10^{-5}$ | $1.72 \times 10^{-7}$ | $1.74 \times 10^{-8}$ |
| CV<br>(Time<br>7946) | 0.0134 | $9.11 \times 10^{-3}$ | $4.28 \times 10^{-17}$ | 0 | $2.62 \times 10^{-3}$ | 0.0385 | 0.0426 | $8.94 \times 10^{-3}$ | $8.94 \times 10^{-3}$ |
| #<br>Interactions | 1 | 2 | 1 | 5 | 1 | 1 | 1 | 2 | 2 |

## Discussion

Two main conclusions can be drawn from the research presented in this paper. First, the number of other compounds with which a compound interacts does not increase the variability of the compound. Second, there were apparent differences between the stochastic and deterministic models’ results. Initially we suspected that the differing behavior between the two models resulted from reactions occuring more times in the stochastic model. However, further investigation showed that increasing the number of reactions in the deterministic model has no significant effect on the variables’ behavior (see supplementary figure.5). This suggests another cause, instead of the number of reactions being the cause for the different behavior between models, the differences could arise from the differing number of reactions between species. In other words, the variables not occurring the same number of times as each other in the stochastic model is the cause of the differences.

Based on the research here, it is impossible to say if either model is better than the other. However, we can say that they provide different insights; thus, each likely has its own strengths and weaknesses. The reactions not occurring a set number of times in the stochastic model seemed to be central to the different results between the two models.

## Acknowledgments

I would like to acknowledge the support and suggestions provided by Doctor Christopher McFarland, Doctor Peter Thomas, and Doctor Ann Harris from Case Western Reserve University along with the Functional Genomics Training program. I would also like to acknowledge Professor Christopher Leary and Professor Gregg Hartvigsen for their advice regarding the deterministic model. Meaghan Parks was supported by NIH grants T32-GM135081 and R00-CA226506.

## Supporting Information Captions

SFigure 1 Graph showing stochastic vs deterministic behavior of amyloid beta inside the neuron

Sfigure 2 Graph showing stochastic vs deterministic behavior of NFTs inside the neuron

Sfigure 3 Graph showing stochastic vs deterministic behavior of NFTs outside the neuron

Sfigure 4 Stochastic vs deterministic behavior of immune cells.

Sfigure 5 Results from the deterministic model with various time steps

## Notes

### Competing Interest Statement

The authors have declared no competing interest.

