## Supplementary figures and images for "Stochastic model of Alzheimer’s Disease progression using two-state Markov chains"

### Supplemental Figure 1

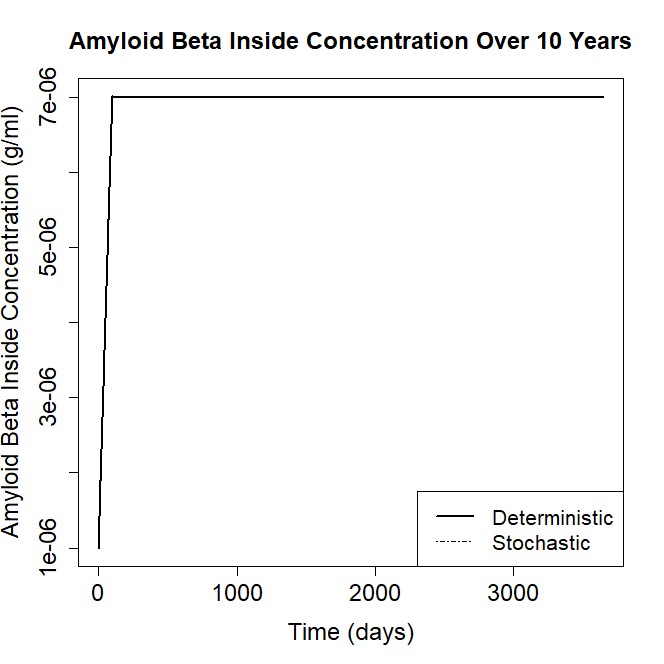

### Supplemental Figure 2

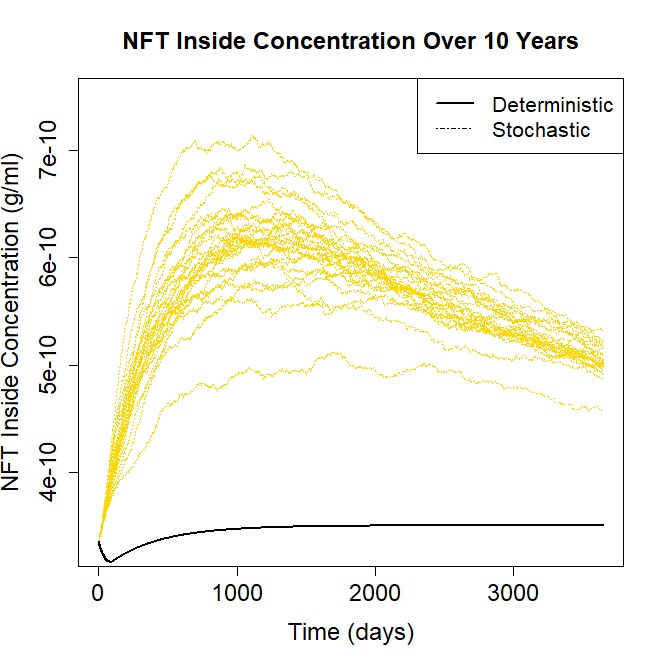

### Supplemental Figure 3

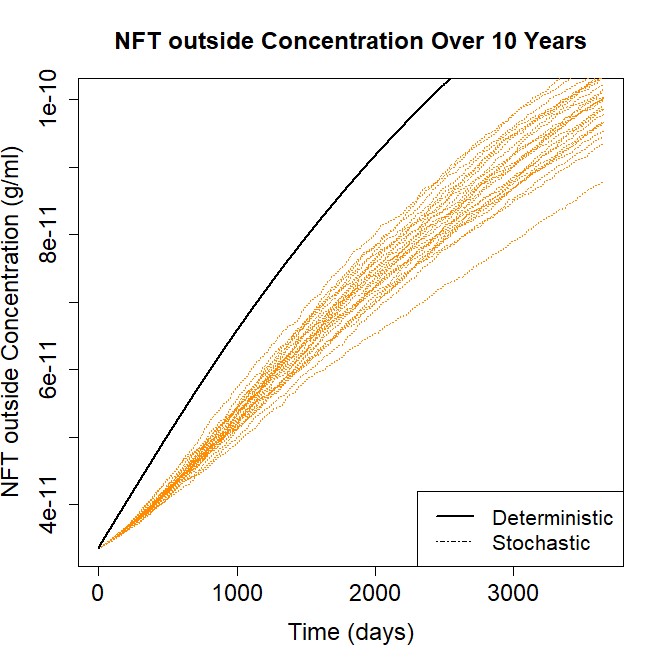

### Supplemental Figure 4

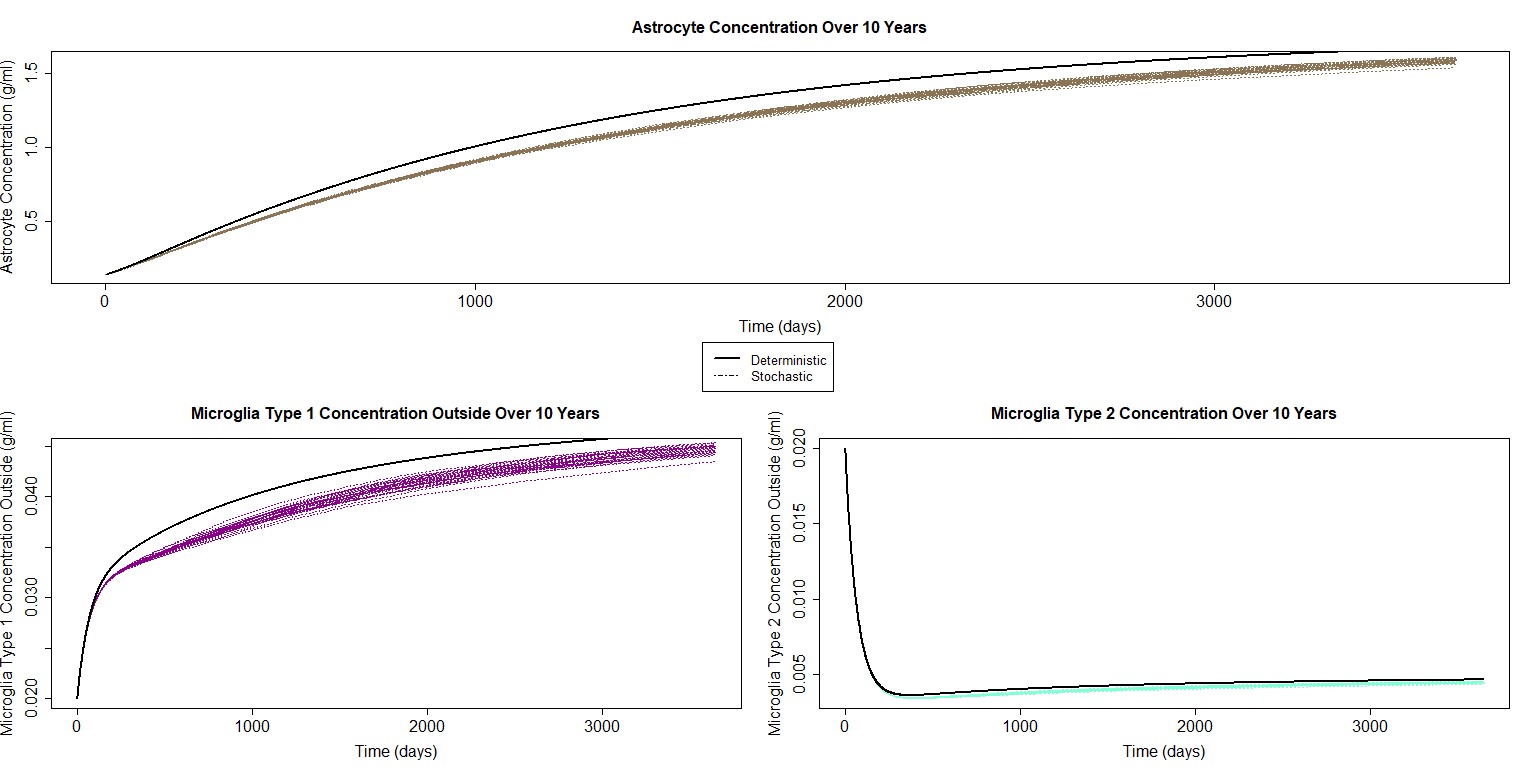

### Supplemental Figure 5

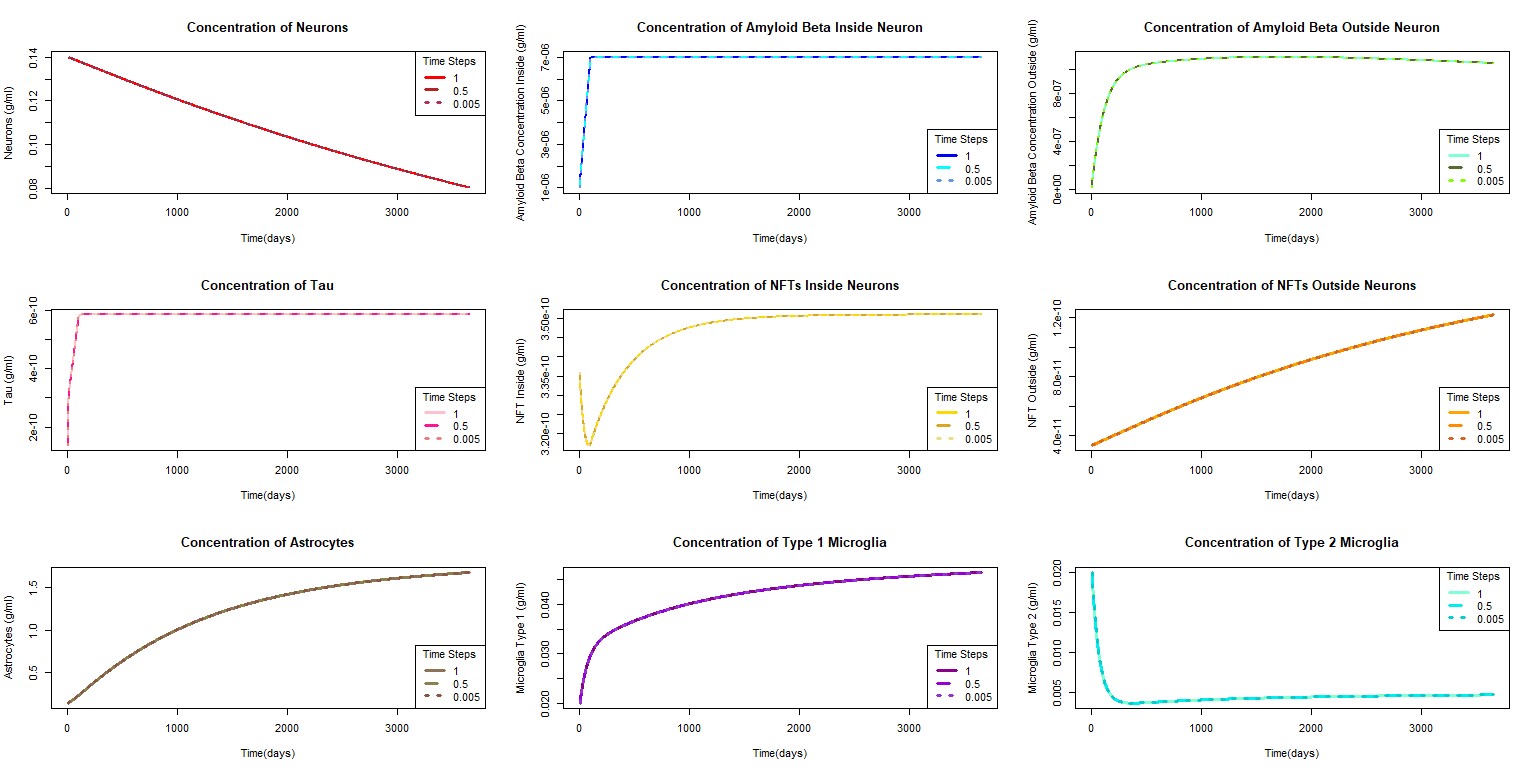
